# The protective effect of club cell secretory protein (CC-16) on COPD risk and progression: a Mendelian randomisation study

**DOI:** 10.1101/2019.12.20.885384

**Authors:** Stephen Milne, Xuan Li, Ana I Hernandez Cordero, Chen Xi Yang, Michael H Cho, Terri H Beaty, Ingo Ruczinski, Nadia N Hansel, Yohan Bossé, Corry-Anke Brandsma, Don D Sin, Ma’en Obeidat

## Abstract

**Background:** There are currently no robust biomarkers of chronic obstructive pulmonary disease (COPD) risk or progression. Club cell secretory protein-16 (CC-16) is associated with the clinical expression of COPD. We aimed to determine if there is a causal effect of serum CC-16 level on COPD risk and/or progression using Mendelian randomisation (MR) analysis.

**Methods:** We performed a genome-wide association meta-analysis for serum CC-16 in two COPD cohorts (Lung Health Study [LHS], n=3,850 and ECLIPSE, n=1,702). We then used the CC-16-associated single-nucleotide polymorphisms (SNPs) in MR analysis to estimate the causal effect of serum CC-16 on COPD risk (International COPD Genetics Consortium/UK-Biobank dataset; n=35,735 cases, n=222,076 controls) and progression (change in forced expiratory volume in 1 s [FEV_1_] in LHS and ECLIPSE). We also determined the associations between SNPs associated with CC-16 and gene expression using n=1,111 lung tissue samples from the Lung eQTL Study.

**Results:** We identified 7 SNPs independently associated (p<5×10^−8^) with serum CC-16 levels; 6 of these were novel. MR analysis suggested a protective causal effect of increased serum CC-16 on COPD risk (p=0.008) and progression (LHS only, p=0.02). Five of the SNPs were also associated with gene expression in lung tissue, including that of the CC-16-encoding gene *SCGB1A1* (false discovery rate<0.1).

**Conclusion:** We have identified several novel genetic variants associated with serum CC-16 level in COPD cohorts. These genetic associations suggest a potential causal effect of serum CC-16 on COPD risk and progression. Further investigation of CC-16 as a biomarker or therapeutic target in COPD is warranted.

**KEY MESSAGES:** **What is the key question?**

Can genetics help uncover a causal effect of serum CC-16 level on COPD risk and/or progression?

**What is the bottom line?**

There is a protective effect of genetically-increased serum CC-16 on both COPD risk and progression (as measured by change in FEV_1_ over time), which may be due to increased expression of the CC-16-encoding gene *SCGB1A1* in the lung.

**Why read on?**

This is the first study to demonstrate a possible causal effect of serum CC-16 in people with COPD, and highlights the potential for CC-16 as a biomarker or therapeutic target.

## INTRODUCTION

Chronic obstructive pulmonary disease (COPD) is expected to be the third leading cause of death worldwide by 2030.[1] Currently, there are no acceptable biomarkers for predicting the risk of COPD or its progression. If robust biomarkers and disease-modifying therapies are to be developed, a better understanding of the causal mechanisms underlying the risk of COPD, and its progression over time, is critical.

Club Cell Secretory Protein 16 (CC-16) is a 16 kDa pneumoprotein produced predominantly by club cells in the airway epithelium, where it appears to have a protective, anti-inflammatory effect.[2–4] Reduced CC-16 levels are associated with the clinical expression of COPD: serum, sputum, and bronchoalveolar lavage fluid (BALF) CC-16 concentrations decrease with increasing disease severity;[5–8] and reduced serum CC-16 is associated with faster lung function decline.[9, 10] In a mouse model of cigarette smoke exposure, CC-16^−/−^ knockout mice showed greater COPD-like lung changes than wild type mice.[3] However, this finding was not replicated by other investigators.[10] While this discrepancy may be explained by differences in experimental design (including the duration and dose of smoke exposure), it is clear that further efforts to establish a causal role for reduced CC-16 in COPD pathogenesis are required.

Mendelian randomisation (MR) analysis is a promising approach to explore causality in humans by relating genetic variants to disease outcomes via a biological risk factor (e.g. protein levels).[11] Due to the random allocation of alleles at meiosis, and under the assumption that the genetic variants exert their effects on disease outcomes only *via* the risk factor of interest, MR analysis is relatively resistant to confounding. Furthermore, since the genomic contribution to the risk factor is constant over a lifetime, MR analysis is relatively resistant to bias due to reverse causation (where the disease itself influences the protein level). For these reasons, MR analysis is particularly attractive for establishing a potential causal role for biological factors in complex diseases such as COPD. We have previously demonstrated a potential causal effect of low surfactant protein D (SPD) in increasing COPD risk and progression using the MR framework.[12] Other molecules associated with COPD outcomes in epidemiological studies, such as C-reactive protein,[13] interleukin-6[14] and blood eosinophil count,[15] do not yield evidence of causality when subjected to analysis by MR. Therefore, MR analysis may be useful in identifying the most promising biological pathways for biomarker and/or drug development in COPD.

In this study, we aimed to determine if there is any causal effect of reduced serum CC-16 level on the risk of COPD and the rate of its progression. We performed the largest genome-wide association study (GWAS) for serum CC-16 to date, and uncovered known, as well as many novel, genetic loci associated with CC-16 levels. These loci were then utilised in a MR analysis to show that genetically-increased serum CC-16 levels reduce the risk of having COPD and slow the rate of lung function decline in people with established COPD. Additionally, we determined that at least some of these effects may be due to increased lung tissue expression of the CC-16-encoding gene, secretoglobin family 1A member 1 (*SCGB1A1*). Our findings reinforce the importance of CC-16 in COPD pathogenesis, and its potential as a biomarker and therapeutic target.

## METHODS

### Study overview

The study workflow is outlined in Figure 1. We first examined the relationship between serum CC-16 levels and changes in lung function over time. Next, we identified genetic variants (single-nucleotide polymorphisms [SNPs]) independently associated with serum CC-16 levels in a GWAS. We then determined the associations between the SNPs associated with CC-16 levels and two COPD outcomes: 1) the presence of COPD in a case-control dataset (“COPD risk”); and 2) the rate of change in lung function (“COPD progression”). Following this, we performed MR analyses for COPD risk and progression using the SNPs associated with CC-16 levels as instrumental variables and serum CC-16 level as the risk factor. Finally, we determined which of the CC-16-associated SNPs were also quantitatively associated with gene expression (mRNA levels) in lung tissue. Access to data from each of the studies examined was granted by the respective governing committees. Participant consent and institutional ethics approval was given for each cohort study. For the present analysis, participants were not identified nor contacted, and further institutional ethics approval was not required.

**Figure 1:**
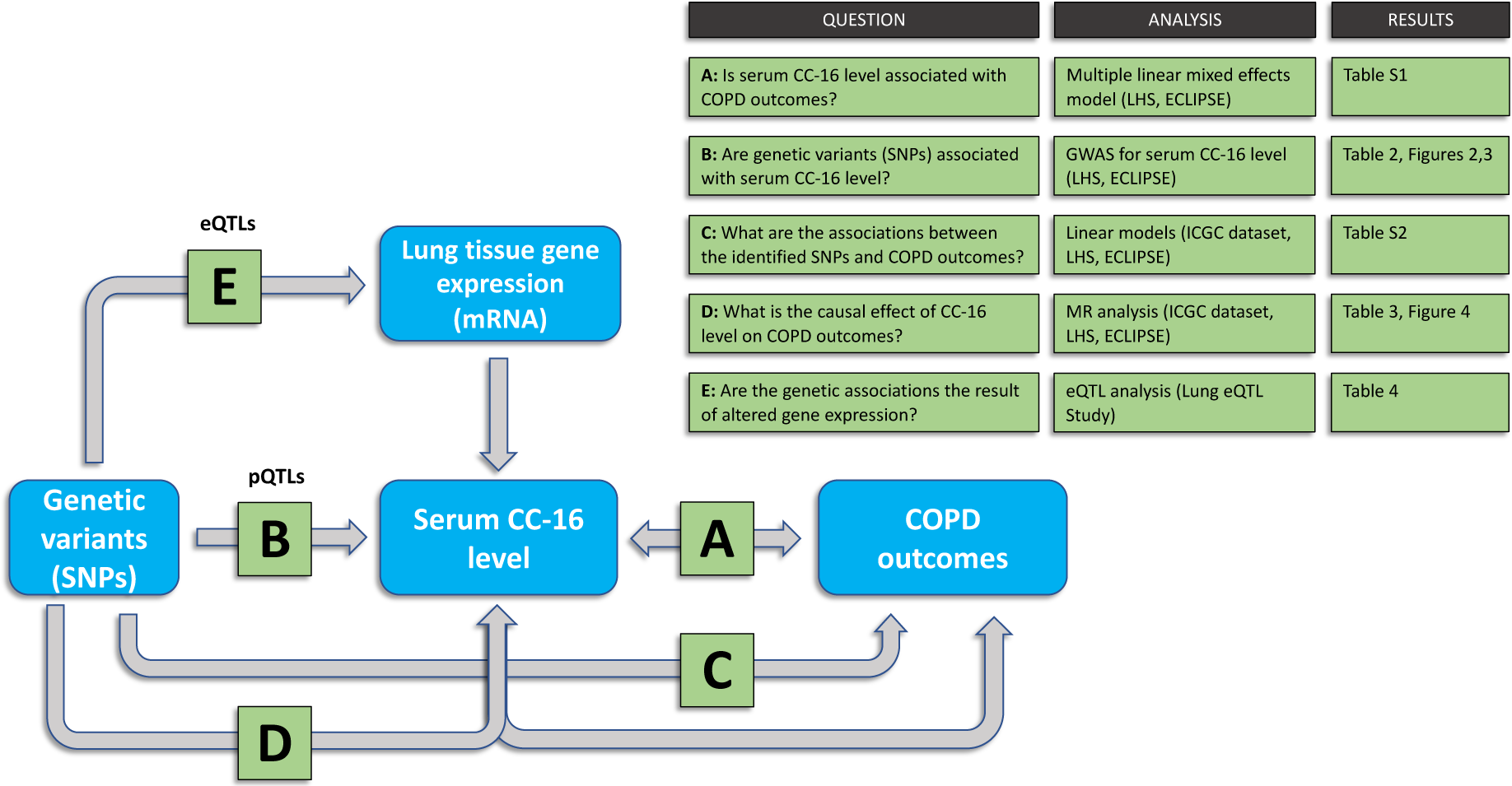
Study workflow. Under the Mendelian randomisation (MR) framework, genetic variants are associated with the phenotype via the risk factor/exposure. MR analysis therefore attempts to estimate the causal effect of the exposure. The relationship may be mediated by altered gene expression. CC-16, club cell secretory protein-16; SNP, single-nucleotide polymorphism; COPD, chronic obstructive pulmonary disease; LHS, Lung Health Study; ICGC dataset, meta-analysis of International COPD Genetics Consortium and UK Biobank COPD cases and non-COPD controls; eQTL, expression quantitative trait locus; pQTL, protein quantitative trait locus.

### Description of study cohorts

Detailed descriptions of the included studies are presented in the Supplementary Methods. Brief summaries are provided below.

#### Lung Health Study (LHS) cohort:[16]

LHS was a longitudinal study examining the effects of smoking cessation intervention and regular inhaled bronchodilator on the rate of lung function decline in 5,887 smokers aged 35-60 years with mild-moderate COPD. Lung function was measured annually for the first 5 years and at year 11. At Year 5, the investigators collected blood samples for measurement of blood biomarkers and genotyping (see Supplementary Methods).

#### Evaluation of COPD Longitudinally to Identify Predictive Surrogate Endpoints (ECLIPSE) cohort:[17]

ECLIPSE was a longitudinal study of 2,746 current or former smokers aged 40-75 years, who were followed for 3 years. COPD cases were defined by spirometry at baseline. Lung function measurements were performed at baseline, 3 months, 6 months, and every 6 months thereafter. Blood was collected at baseline for biomarker analysis and genotyping (see Supplementary Methods).

#### 2019 International COPD Genetics Consortium (ICGC)/UK Biobank dataset:[18]

ICGC is an international collaboration that pooled genome-wide association data of COPD cohort, case-control, and general population studies that all performed spirometry. A recent meta-analysis of 35,735 COPD cases and 222,076 non-COPD controls of European ancestry from the ICGC cohorts and the UK Biobank (a general population cohort of 502,682 volunteers) identified genetic variants with a genome-wide significant association with risk of COPD. A detailed description of this meta-analysis is provided in the Supplementary Methods, but for simplicity this will be referred to as the “ICGC dataset”. For our analysis, we used the summary statistics resulting from the meta-analysis.

#### Lung eQTL Study:[19]

This study was a meta-analysis of the association between SNPs and gene expression in non-tumour lung tissue from 1,111 volunteers (some of which had COPD) undergoing lung resection at three different centres, adjusted for age, sex, and smoking status.

### Serum CC-16 quantification

Serum CC-16 quantification in the LHS and ECLIPSE has been described previously[8, 10] (see Supplementary Methods). We transformed serum CC-16 concentrations by their natural logarithm to mitigate the influence of extreme outliers and approximate normality in our analysis.

### Association between serum CC-16 level and COPD progression

We tested for associations between serum CC-16 concentration and COPD progression (annual change in forced expiratory volume in 1 s [FEV_1_]) in the the LHS and ECLIPSE using a multiple linear mixed effects (LME) model adjusting for age, sex, body mass index (BMI), smoking status, baseline FEV_1_, and their factor by time interactions as fixed effects, and time as a random effect (Analysis A in Figure 1). We then combined the results of these two studies in a meta-analysis.

### Genome-wide association study (GWAS) for serum CC-16 concentration

The full details of the GWAS for serum CC-16 level are outlined in the Supplementary Methods. Briefly, we identified SNPs associated with serum CC-16 concentration (protein quantitative trait loci [pQTLs], Analysis B in Figure 1) in the LHS and ECLIPSE and combined the results in an inverse variance weighted (IVW) fixed-effects meta-analysis. We identified independently-associated SNPs using conditional analysis within each 2 Mb gene region.[20] From this, we retained only SNPs having independent, genome-wide significant (p<5×10^−8^) associations with CC-16 levels.

### Associations between CC-16 pQTLs and COPD outcomes

For COPD risk, we extracted summary statistics for each CC-16 pQTL’s association with the presence of COPD in the ICGC dataset. For COPD progression, we tested serum CC-16 pQTLs for possible association with changes in FEV_1_ over time in the LHS and ECLIPSE cohorts separately, using the LME model described previously, with the addition of the first 5 genetic principal components as covariates (Analysis C in Figure 1).

### Mendelian randomisation (MR) analysis

To estimate the causal effect of CC-16 concentration on COPD outcomes (Analysis D in Figure 1), we related the CC-16 pQTL per-allele effects on serum CC-16 levels to their effects on COPD outcomes using an IVW MR model with the CC-16 pQTLs as instrumental variables, with nominal significance set at p<0.05 (see Supplementary Methods for description of the model). To test for violations of MR assumptions,[21] we tested for heterogeneity by Cochran’s Q statistic, and for evidence of directional pleiotropy by both MR-Egger[22] (intercept term) and MR-PRESSO (“global test”),[23] with nominal significance of each test set at p<0.05 (see Supplementary Methods).

### Lung tissue gene expression analysis

To determine if the serum CC-16 pQTLs have any effects on quantitative gene expression in lung tissue, we tested for their associations (at false discovery rate [FDR]<0.1) with mRNA transcript levels using data from the Lung eQTL Study[19] (Analysis E in Figure 1) (See Supplementary Methos). For this analysis, we considered only *cis*-eQTLs, within a region 1 Mb either side of the sentinel SNP.

### Statistical software

We performed all analyses in R (version 3.4.0; www.r-project.org).[24]

## RESULTS

### Serum CC-16 level is associated with COPD progression

Table 1 summarises the demographics, lung function, and CC-16 concentrations in the two biomarker cohorts (LHS and ECLIPSE). Serum CC-16 level was significantly associated with changes in FEV_1_ in the LHS (beta=2.43, p=0.009) but not in ECLIPSE (beta=6.25, p=0.11) (Supplementary Results Table S1). When combined by meta-analysis, this translated to a 2.64 mL/year slower decline in FEV_1_ per 1 ln(ng/mL) increase in serum CC-16 concentration (p=0.004).

**Table 1:**
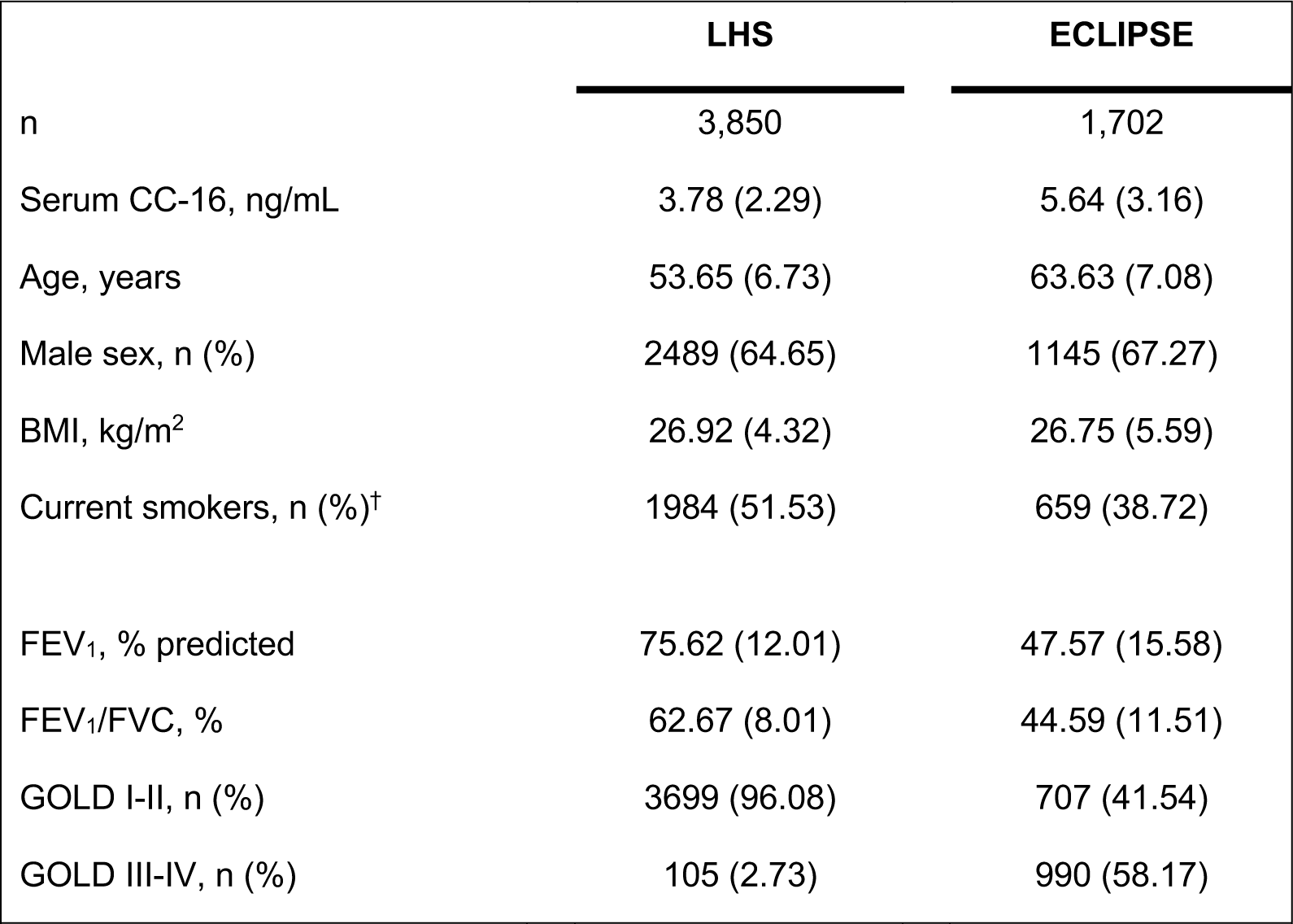
Characteristics of the biomarker cohorts for serum CC-16 genome-wide association study (GWAS). All values are mean (SD) unless otherwise stated. ^†^smoking status determined from study data (see Supplementary Methods). CC-16, club cell secretory protein-16; LHS, Lung Health Study; BMI, body mass index; FEV_1_, forced expiratory volume in 1 s; FVC, forced vital capacity; GOLD-I-II, FEV_1_ ≥50 percent predicted; GOLD III-IV, FEV_1_ <50 percent predicted.

|  | LHS | ECLIPSE |
| --- | --- | --- |
| n | 3,850 | 1,702 |
| Serum CC-16, ng/mL | 3.78 (2.29) | 5.64 (3.16) |
| Age, years | 53.65 (6.73) | 63.63 (7.08) |
| Male sex, n (%) | 2489 (64.65) | 1145 (67.27) |
| BMI, kg/m<sup>2</sup> | 26.92 (4.32) | 26.75 (5.59) |
| Current smokers, n (%)<sup>†</sup> | 1984 (51.53) | 659 (38.72) |
| FEV<sub>1</sub>, % predicted | 75.62 (12.01) | 47.57 (15.58) |
| FEV<sub>1</sub>/FVC, % | 62.67 (8.01) | 44.59 (11.51) |
| GOLD I-II, n (%) | 3699 (96.08) | 707 (41.54) |
| GOLD III-IV, n (%) | 105 (2.73) | 990 (58.17) |

### Multiple novel loci are associated with serum CC-16 level (pQTLs)

We included 7,312,348 SNPs in a total of 5,552 individuals (n= 3,850 in LHS, n= 1,702 in ECLIPSE) in the GWAS. A total of 7 pQTLs showed independent, genome-wide significant association with serum CC-16 level in the meta-analysis (Figures 2 [Manhattan plot] and 3 [gene region plots]; Table 2). Quantile-quantile plots showed deviation from the expected distribution at low p values (Supplementary Results Figure S3), indicating a strong association. Of the loci identified in this meta-analysis, only SNP rs3741240 on chromosome 11 (an intronic variant in the 5’ untranslated region of *SCGB1A1*, the gene which encodes the CC-16 protein) had been previously identified in a GWAS for serum CC-16 (performed in the ECLIPSE cohort).[25] The remaining 6 loci are novel pQTLs for serum CC-16: SNP rs4971100 (chromosome 1) is an intronic variant in the tripartite motif containing 46 (*TRIM46*) gene; rs1515498 (chromosome 3) is an intronic variant in the tumour protein p63 (*TP63*) gene; rs37002 is in an intergenic region on the short arm of chromosome 5 (p15.3); rs11032840 is an intronic variant in an uncharacterised region on the short arm of chromosome 11; rs11231085 (chromosome 11) is an intronic variant in the *SCGB1A1* gene; and rs7962469 (chromosome 12) is an intronic variant in the activin A receptor type 1B (*ACVR1B*) gene.

**Table 2:** Genome-wide association studies (GWAS) for serum CC-16 level. Separate GWAS for each cohort, combined by inverse variance weighted meta-analysis. *p value less than genome-wide significant cut-off of 5×10^−8^. LHS, Lung Health Study; ECLIPSE, Evaluation of COPD Longitudinally to Identify Predictive Surrogate Endpoints; SNP, single nucleotide polymorphism; rsID, reference SNP cluster identifier; Chr, chromosome; MAF, minor allele frequency; *TRIM46*, tripartite motif containing 46; *TP63*, tumour protein p63; *SCGB1A1*, secretoglobin family 1A member 1; *ACVR1B*, activin receptor 1B.

| SNP rsID | Chr | Position | Effect allele | Alternate allele | Nearby gene | LHS |  |  |  | ECLIPSE |  |  |  | Meta-analysis |  |
| --- | --- | --- | --- | --- | --- | --- | --- | --- | --- | --- | --- | --- | --- | --- | --- |
|  |  |  |  |  |  | MAF | Allele effect | Allele effect | % variance explained | MAF | Allele effect | Allele effect | % variance explained | Allele effect | Allele effect |
| | | | | | | | $\beta$ | p | | | $\beta$ | p | | $\beta$ | p |
| rs4971100 | 1 | 155155731 | A | G | *TRIM46* | 0.42 | 0.08 | $5.69 \times 10^{-9*}$ | 0.58% | 0.41 | 0.06 | $5.66 \times 10^{-4}$ | 0.60% | 0.07 | $1.55 \times 10^{-11*}$ |
| rs1515498 | 3 | 189508302 | A | G | *TP63* | 0.36 | 0.07 | $1.38 \times 10^{-7}$ | 0.58% | 0.34 | 0.06 | $8.62 \times 10^{-4}$ | 0.43% | 0.07 | $4.94 \times 10^{-10*}$ |
| rs37002 | 5 | 1356944 | C | T | - | 0.49 | 0.05 | $6.95 \times 10^{-4}$ | 0.28% | 0.49 | 0.08 | $3.92 \times 10^{-6}$ | 1.14% | 0.06 | $3.15 \times 10^{-8*}$ |
| rs11032840 | 11 | 34779464 | G | T | - | 0.40 | 0.06 | $3.72 \times 10^{-6}$ | 0.38% | 0.40 | 0.10 | $2.75 \times 10^{-8*}$ | 1.29% | 0.08 | $1.44 \times 10^{-12*}$ |
| rs3741240 | 11 | 62186542 | G | A | *SCGB1A1* | 0.36 | 0.18 | $1.1 \times 10^{-37*}$ | 3.95% | 0.35 | 0.17 | $4.20 \times 10^{-22*}$ | 4.99% | 0.18 | $2.08 \times 10^{-59*}$ |
| rs11231085 | 11 | 62190448 | G | C | *SCGB1A1* | 0.35 | 0.18 | $1.94 \times 10^{-39*}$ | 3.66% | 0.35 | 0.16 | $6.03 \times 10^{-18*}$ | 4.28% | 0.17 | $9.13 \times 10^{-57*}$ |
| rs7962469 | 12 | 52348259 | A | G | *ACVR1B* | 0.34 | 0.06 | $3.87 \times 10^{-5}$ | 0.38% | 0.34 | 0.07 | $1.25 \times 10^{-4}$ | 0.47% | 0.06 | $1.99 \times 10^{-8*}$ |

**Figure 2:**
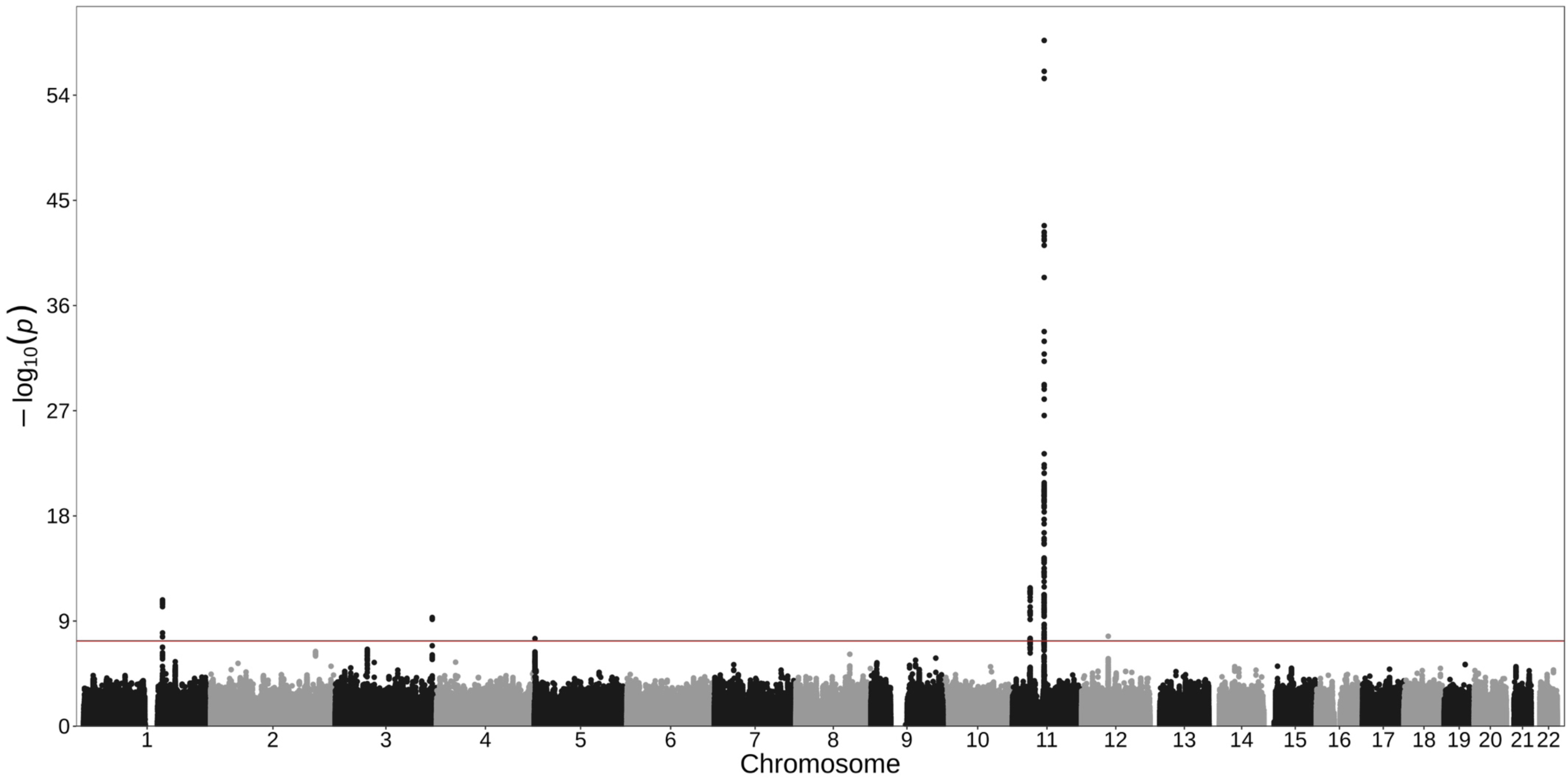
Manhattan plot for serum CC-16 genome-wide association study (GWAS). Meta-analysis of Lung Health Study and ECLIPSE study GWAS. GWAS p values (– log_10_ scale) (Y axis) versus single nucleotide polymorphism positions across 22 chromosomes (X axis). Horizontal red line represents the genome-wide significance cut-off of 5×10^−8^. CC-16, club cell secretory protein-16.

### Serum CC-16 pQTLs are associated with COPD risk and progression

The effect estimates for individual serum CC-16 pQTLs on COPD outcomes are shown in the Supplementary Results (Table S2). One pQTL (rs7962469 on chromosome 12) was significantly associated with COPD risk in the ICGC dataset (p=1.2×10^−3^). Two pQTLs (rs3741240 and rs11231085, both on chromosome 11) were significantly associated with change in FEV_1_ in the LHS (p=0.047 and p=5.4×10^−4^, respectively). rs3741240 was also significantly associated with change in FEV_1_ in ECLIPSE (p=0.02).

### Genetically-determined serum CC-16 level is related to COPD risk and progression

The IVW MR estimate for COPD risk in the ICGC dataset was significant (beta=–0.11, p=0.008), suggesting a protective effect of genetically-increased serum CC-16 level on the risk of COPD (Table 3, Figure 4A). Tests for violation of MR assumptions were not statistically significant (Table 3; Supplementary Results Figure S4).

**Table 3:** Mendelian randomisation (MR) analysis for effect of genetically-determined serum CC-16 level on COPD outcomes. Primary MR analysis by inverse variance weighted (IVW) linear regression, with Cochran’s Q test for heterogeneity, MR-Egger and MR-PRESSO tests for directional pleiotropy. *p<0.05. ICGC, International COPD Genetics Consortium; LHS, Lung Health Study; FEV_1_, forced expiratory volume in 1 s; SE, standard error.

|  |  | IVW MR |  |  | MR-Egger |  |  |  | MR-PRESSO |
| --- | --- | --- | --- | --- | --- | --- | --- | --- | --- |
| Cohort | Outcome variable | Estimate<br>β (SE) | Estimate<br>p | Cochran's Q<br>p | Estimate<br>β (SE) | Estimate<br>p | Cochran's Q<br>p | Intercept<br>p | Global test<br>p |
| COPD risk |  |  |  |  |  |  |  |  |  |
| ICGC dataset | COPD (yes/no) | -0.11 (0.04) | 0.008* | 0.16 | -0.03 (0.11) | 0.75 | 0.13 | 0.45 | 0.29 |
| COPD Progression |  |  |  |  |  |  |  |  |  |
| LHS | Change in FEV<sub>1</sub><br>(mL/year) | 7.40 (3.28) | 0.02* | 0.05 | 22.18 (8.50) | 0.009* | 0.17 | 0.05 | 0.08 |
| ECLIPSE | Change in FEV<sub>1</sub><br>(mL/year) | 19.14 (12.59) | 0.13 | 0.21 | - | - | - | - | 0.27 |

**Figure 3:**
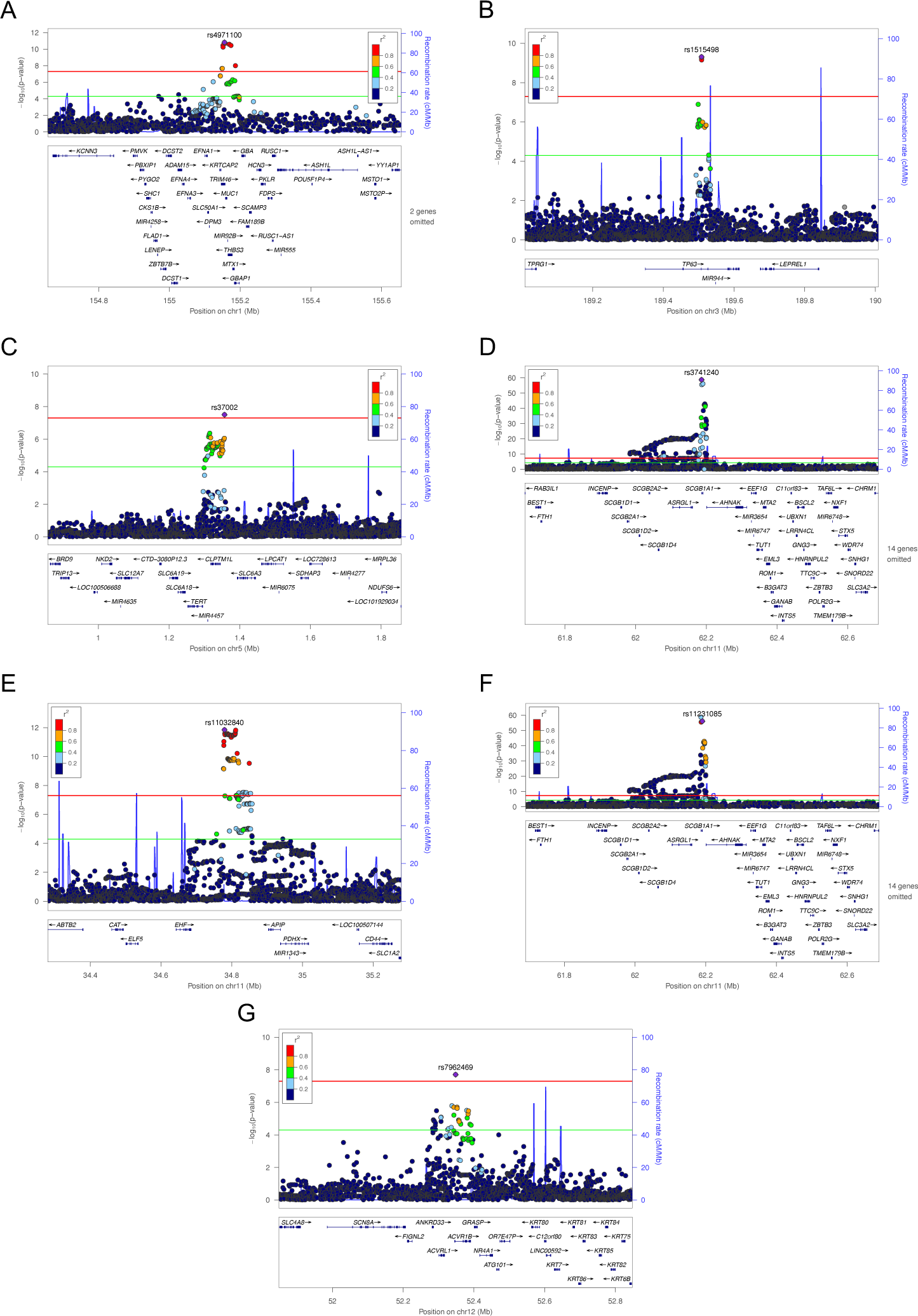
Gene region plots for serum CC-16 genome-wide association study (GWAS). Meta-analysis of Lung Health Study and ECLIPSE study GWAS. GWAS p values (–log_10_ scale) (Y axes) versus single-nucleotide polymorphism (SNP) genomic position (X axes) on (A) chromosome 1, (B) chromosome 3, (C) chromosome 5, (D-F) three independently-associated regions on chromosome 11, and (G) chromosome 12. Horizontal red line represents the genome-wide significance cut-off of 5×10^−8^. Horizontal green line represents p-value cut-off of 5×10^−5^. Gene names and their coordinates are labeled. The sentinel (most significant) SNP for each region is annotated by rs identifier. r^2^, linkage disequilibrium with the sentinel SNP.

**Figure 4:**
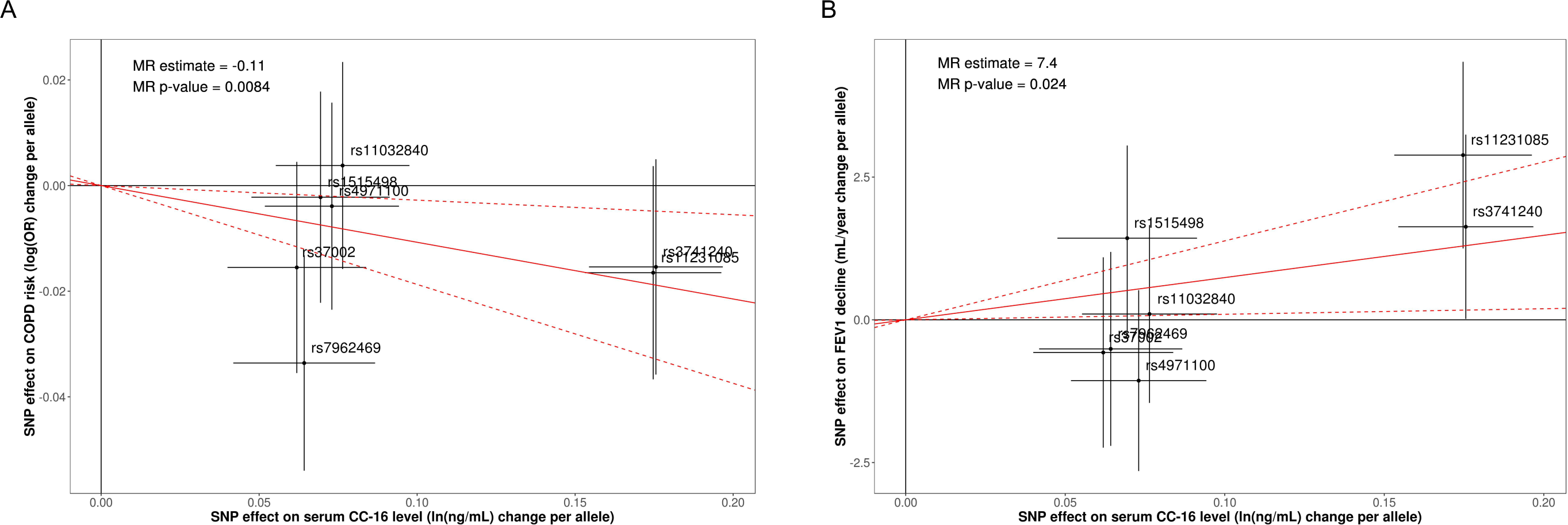
Mendelian randomisation (MR) plots. Inverse variance weighted regression model, adjusted for linkage disequilibrium between single-nucleotide polymorphisms (SNPs), intercept constrained to zero. The model relates the per-allele effects of the SNPs on serum CC-16 level (X axes) to their per-allele effects on (A) risk of chronic obstructive pulmonary disease (COPD) in the ICGC dataset, and (B) change in forced expiratory volume in 1 s (FEV_1_) over time in the LHS (Y axes). Red line represents the estimated effect. Error bars represent 95% confidence intervals. Individual SNPs are annotated by their rs identifier. CC-16, club cell secretory protein-16; OR, odds ratio.

The IVW MR estimate for change in FEV_1_ in the LHS was statistically significant (beta=7.4, p=0.02), suggesting a protective effect of genetically-increased serum CC-16 on COPD progression (Table 3, Figure 4B). Again, tests for violations of MR assumptions were not statistically significant (Table 3; Supplementary Results Figure S5). The IVW MR estimate for change in FEV_1_ in the ECLIPSE study was not significant (Table 3; Supplementary Results Figure S6).

### Multiple CC-16 pQTLs affect lung tissue gene expression

Four of the pQTLs for serum CC-16 level were also lung eQTLs (Table 4). SNPs rs3741240 and rs11231085 on chromosome 11 were associated with levels of *SCGB1A1* mRNA (p=4.13×10^−12^ and p=3.82×10^−7^, respectively), with allele effects in the same direction as their effects on serum CC-16 level. SNP rs4971100 (chromosome 1) was associated with mRNA levels for the glucosylceramidase beta (*GBA*) gene (p=4.58×10^−8^) in lung tissue. SNP rs7962469 (chromosome 12) was associated with *ACVR1B* gene expression (p=3.31×10^−116^) with the allele effect in the same direction as its effect on serum CC-16 level.

**Table 4:** Lung tissue expression quantitative trait loci (eQTL) analysis. Expression data from the Lung eQTL Study. SNP, single nucleotide polymorphism; rsID, reference SNP cluster identifier; Chr, chromosome; *GBA*, glucosylceramidase beta; *SCGB1A1*, secretoglobin family 1A member 1; *ACVR1B*, activin receptor 1B.

| SNP rsID | Chr | Position | Effect allele | Alternate allele | Probe set | Gene | Allele effect $\beta$ | Allele effect $p$ |
| --- | --- | --- | --- | --- | --- | --- | --- | --- |
| rs4971100 | 1 | 155155731 | A | G | 100300160_TGI_at | *GBA* | -0.23 | $4.58 \times 10^{-8}$ |
| rs37002 | 5 | 1356944 | C | T | 100307694_TGI_at | Unannotated | 0.93 | $7.47 \times 10^{-161}$ |
| rs3741240 | 11 | 62186542 | G | A | 100143118_TGI_at | *SCGB1A1* | 0.31 | $4.13 \times 10^{-12}$ |
| rs11231085 | 11 | 62190448 | G | C | 100143118_TGI_at | *SCGB1A1* | 0.22 | $3.82 \times 10^{-7}$ |
| rs7962469 | 12 | 52348259 | A | G | 100127400_TGI_at | *ACVR1B* | 0.82 | $3.31 \times 10^{-116}$ |

## DISCUSSION

By performing the largest GWAS-pQTL analysis for serum-CC16 levels to date (n=5,552), we have identified many novel distal loci associated with serum CC-16 level. Furthermore, using the MR framework, we have demonstrated a potential protective effect of genetically-increased serum CC-16 level on both COPD risk and disease progression. Our lung tissue eQTL analysis suggests that genetically-increased serum CC-16 levels may be due to increased lung tissue gene expression at the CC-16-encoding gene *SCGB1A1*. Our findings expand upon the existing body of literature on the biology of CC-16 and support a possible causal role for CC-16 in COPD pathogenesis.

We identified genetic variants independently associated with serum CC-16 level (pQTLs) in 6 different loci across 5 chromosomes. The most significantly-associated SNP was rs3741240 (p=2.08×10-59 in the GWAS meta-analysis), which is in a non-coding region of the CC-16 gene *SCGB1A1* on chromosome 11. This SNP was also the top association in the only previously-published GWAS for serum CC-16 level.[25] In the same gene region, we have identified a second, independently-associated SNP rs11231085 by conditional analysis (p=9.13×10-57 in the GWAS meta-analysis). These results reflect the increased statistical power of GWAS meta-analysis to detect associated genetic variants.

To our knowledge, this is the first study to demonstrate that variants in the *SCGB1A1* gene are associated with lung function decline. The SNP rs3741240 has been previously associated with both airway hyperresponsiveness[26] and asthma,[27] but was not associated with COPD outcomes in the previously-published ECLIPSE GWAS.[25] The SNP rs11231085 has not previously been associated with lung function or lung disease. Our lung tissue eQTL analysis suggests that genetically-increased serum CC-16 levels may be due to increased lung tissue expression of the *SCGB1A1* gene. The precise mechanisms by which *SCGB1A1* gene expression and CC-16 protein production support lung function are unclear.

The remaining genetic variants we found to be associated with serum CC-16 levels were in loci distal to the *SCGB1A1* gene, and each of these associations were novel. SNP rs7962469 on chromosome 12 has been previously associated with risk of lung disease. This SNP is an intronic variant in the *ACVR1B* gene, which codes for a member of the transforming growth factor-beta (TGF-β) receptor superfamily also known as activin-like kinase-4 (ALK4). We found this variant was also associated with lung tissue expression of *ACVR1B* (i.e. it is an eQTL), although how expression of the ALK4 receptor is related to serum levels of CC-16 remains unclear. Members of the TGF-β superfamily, including ALK4, are potent regulators of gene transcription, and the downstream *Smad* signalling pathway has been shown to be dysregulated in COPD.[28] The association between *ACVR1B* variants and serum CC-16 level may therefore be due to modulation of *SCGB1A1* transcription by ALK4 receptor activity. Additionally, ALK4 and *Smad* signalling are involved in the differentiation and proliferation of lung cells including airway epithelial cells.[28] It is therefore possible that *ACVR1B* variants relate to CC-16 levels due to altered numbers of differentiated club cells secreting the CC-16 protein. Interestingly, variants in or near the *ACVR1B* gene region have been associated with lung function[29] and COPD risk.[30, 31] The rs7962469 variant, through its effects on *ACVR1B* expression in lung tissue, has also been causally associated with emphysema distribution on chest imaging in the COPDGene cohort.[30] The link between this gene and CC-16 biology is intriguing, and warrants further investigation.

The links between the other novel SNPs and serum CC-16 level are less clear. SNP rs1515498 on chromosome 3 is an intronic variant in the *TP63* gene, which itself is related to the risk of lung cancer[32, 33] but has also recently been associated with COPD risk in a Taiwanese population.[34] SNP rs4971100 on chromosome 1 is an eQTL for the *GBA* gene in lung tissue, which encodes a cell membrane protein involved in lipid processing. Abnormalities in the *GBA* gene are associated with the lysosomal storage disorder Gaucher’s disease[35] as well as the neurodegenerative disorder Parkinson disease.[36] These distal gene associations highlight the complexity of CC-16 biology.

The association between serum CC-16 levels and lung function is well-documented in observational and epidemiological studies. CC-16 may be critical to lung development.[37] Lower serum CC-16 levels have also been associated with accelerated FEV_1_ decline in early adulthood[38] and increased risk of incident COPD.[37] Despite the consistency of these observations, confounding through reverse causality and environmental exposures remained a possibility. Since MR analysis quantifies only the genomic contribution to serum CC-16 level, it is more resistant to confounding by these factors and argues for a unidirectional association. This is particularly important in light of the disparate effects of cigarette smoke: cumulative chronic exposure is strongly associated with reduced serum CC-16 concentration;[39] yet acute smoke exposure causes a transient increase in serum and commensurate decrease in BALF CC-16 concentration, possibly by disrupting epithelial cell integrity and/or increased vascular permeability.[40] CC-16 appears to have a protective effect on the airway epithelium,[2–4] most likely through regulation of inflammation via the nuclear factor κB (NF-κB) pathway.[3] Genetic and environmental factors that reduce the availability of CC-16 in the lung may therefore increase susceptibility to inflammatory insults, with resultant airway and/or parenchymal changes leading to accelerated lung function decline.

Through MR analysis, we found a significant effect of serum CC-16 level on COPD progression in the LHS. However, we did not find a similar effect in the ECLIPSE study. Notably, the effect estimate in ECLIPSE was similar, suggesting that decreased power due to smaller sample size and shorter follow-up time may explain the discrepancy. Differences in the study populations (LHS, younger cohort with mild-moderate COPD; ECLIPSE, older cohort with more severe COPD and a smaller sample size) may also contribute. Additionally, we did not find a significant relationship between serum CC-16 level and change in FEV_1_ in the ECLIPSE study alone, which is in contrast to previous analyses.[9] This may be due to differences in the models used to determine change in FEV_1_. Nevertheless, the statistically non-significant trend was in the same direction as previously reported effects of serum CC-16 level on change in FEV_1_.[9, 10] Further studies in larger, independent cohorts will be needed to confirm the causal effect of serum CC-16 level on change in FEV_1_.

Although MR analysis suggests a causal effect of serum CC-16 level on COPD outcomes, we took steps to ensure the causal inferences do not violate the fundamental assumptions of MR.[21] These steps included: 1) accounting for linkage disequilibrium structure between the SNPs included in MR analysis, to ensure the instrumental variables were independently associated with serum CC-16 level; 2) testing for heterogeneity, the presence of which would suggest that variability in the SNP associations is greater than would be expected by chance alone and thus may indicate bias; and 3) testing for evidence of directional pleiotropy i.e. one or more of the instrumental variables is associated with the outcome via a risk factor other than serum CC-16, through MR-Egger and MR-PRESSO analyses. When the MR estimates for the effects of serum CC-16 level on both COPD risk and progression were significant, these statistical tests for heterogeneity and directional pleiotropy were not significant, suggesting no major violation of the MR assumptions.

Our results complement previous *in vitro* and animal studies investigating a causal role for CC-16 in COPD pathogenesis. Among the most convincing evidence is the study by Laucho-Contreras and colleagues[3] in which CC-16-/- knockout mice showed signs of accelerated emphysema-like lung changes, airway remodelling, alveolar apoptosis, and increased lung inflammation following cigarette smoke exposure. These effects were attenuated by transfection with a CC-16-producing adenoviral vector. However, these changes were not observed in a similar study by Park and colleagues, despite the use of an identical strain of mouse and a similar experimental design.[10] The reasons for this discrepancy are unclear. Our results therefore contribute important evidence from human studies that CC-16 is implicated in the development and progression of COPD.

## CONCLUSION

Using GWAS and the MR framework, we have provided evidence for a potential causal effect of serum CC-16 level in COPD pathogenesis and lung function decline. These results complement previous attempts to establish a causal role for CC-16 in animal models. Further investigation of serum CC-16 as a biomarker, and its potential as a therapeutic target, in COPD is warranted.

## Supporting information

Supplementary Results

## ACKNOWLEDGEMENTS

This research has been conducted using the UK Biobank Resource. We thank the International COPD Genetics Consortium investigators for providing genetic association summary statistics.

