## Supplementary Methods for "The protective effect of club cell secretory protein (CC-16) on COPD risk and progression: a Mendelian randomisation study"

**AUTHORS:**

Stephen Milne[1,2,3], Xuan Li[1], Ana I Hernandez Cordero[1], Chen Xi Yang[1], Michael Cho[4], Terri H Beaty[5], Ingo Ruczinski[6], Nadia N Hansel[7], Yohan Bossé[8], Corry-Anke Brandsma[9], Don D Sin[1,2], Ma'en Obeidat[1]

1. Centre for Heart Lung Innovation, St Paul's Hospital and University of British Columbia, Vancouver, BC, Canada
2. Division of Respiratory Medicine, Faculty of Medicine, University of British Columbia, Vancouver, BC, Canada
3. Faculty of Medicine and Health, University of Sydney, Sydney, New South Wales, Australia
4. Channing Division of Network Medicine and Division of Pulmonary and Critical Care Medicine, Brigham and Women's Hospital, Boston, MA, USA
5. Department of Epidemiology, Bloomberg School of Public Health, Johns Hopkins University, Baltimore, MD, USA
6. Department of Biostatistics, Bloomberg School of Public Health, Johns Hopkins University, Baltimore, MD, USA
7. Pulmonary and Critical Care Medicine, School of Medicine, Johns Hopkins University, Baltimore, MD, USA
8. Institut universitaire de cardiologie et de pneumologie de Québec, Department of Molecular Medicine, Laval University, Quebec City, Canada
9. University of Groningen Department of Pathology and Medical Biology, University Medical Centre Groningen, Groningen, The Netherlands

**CORRESPONDING AUTHOR:**

Dr Stephen Milne
UBC Centre for Heart Lung Innovation
Rm 166, St Paul's Hospital
1081 Burrard Street,
Vancouver, BC, V6Z 1Y6
CANADA

E:

T: +1 604 806 8346

### 1. DATASETS ANALYSED FOR SERUM CC-16 GENOME-WIDE ASSOCIATION STUDY

#### Description of study populations

*Lung Health Study*

The details of the LHS have been previously published.[1-3] The original LHS ("LHS-I") was a longitudinal study examining the effects of a smoking cessation intervention (including counselling and nicotine replacement therapy) and regular inhaled bronchodilator (ipratropium bromide) on the rate of change in lung function (post-bronchodilator FEV1) in smokers aged 35-60 years with mild-moderate COPD. COPD was defined by post-bronchodilator FEV1 55-90 percent predicted and FEV1/FVC ratio <0.7. A total of 5,887 participants were enrolled in LHS-I.[1] Interviews and spirometry were performed annually for 5 years. An extension of the original LHS, named LHS-III, involved an additional study visit 11 years after enrolment.[2]

In LHS, there was an increase in FEV1 between screening and Year 1, which has been attributed to the effects of smoking cessation.[1] For the remainder of the observation period, there was a general decline in FEV1 over time. Therefore, for the purposes of our analysis, data from Year 1 to 11 were used in the analysis of change in FEV1 over time.

Smoking status was assessed at each visit by self-report, and verified by exhaled carbon monoxide (eCO) and salivary cotinine (sCot) analysis. Data on smoking status was only available to us from LHS-I; we therefore assigned smoking status based on the baseline to year 5 data. We labelled participants as "continuous smokers" (eCO/sCot-verified smoking at all study visits), "sustained quitters" (eCO/sCot-verified non-smoking, plus self-reported number of cigarettes = 0, at every study visit) or "intermittent quitters" (eCO/sCot-verified non-smoking at some but not all study visits, or self-reported smoking despite negative eCO/sCot at some but not all visits). We then used these smoking status labels as covariates in our models for GWAS and lung function decline.

At Year 5, the LHS investigators collected blood samples on 89% of the participants. A description of CC-16 measurement is given below. Genotyping in the LHS was undertaken using the Illumina Human660W-Quad v.1_A BeadChip (Illumina, San Diego, CA, USA), as previously described.[4]

Study protocols in the LHS were approved by the institutional review boards at each trial center, and written informed consent was obtained from each participant

*Evaluation of COPD Longitudinally to Identify Predictive Surrogate End-points (ECLIPSE):*

The details of the ECLIPSE cohort have been previously published.[5] ECLIPSE is a longitudinal study of current or former smokers with ≥10 pack year exposure aged 40-75 years, followed over 3 years. For the purposes of our analysis, we analysed only Caucasian COPD cases defined by post-bronchodilator FEV1 <80 percent predicted and FEV1/FVC ratio ≤0.7. Interviews and spirometry were performed at baseline, 3 months, 6 months and every 6 months after. An increase in mean FEV1 was observed between enrolment and Visit 1 (3 months). Therefore, we used only data from Visit 1 to Visit 7 (3 years) to determine change in FEV1 over time.

Smoking status was assessed at each study visit by interview, and verified by eCO. However, only smoking status from the baseline visit was available to us. We labelled participants as “smoker” (positive self-reported smoking or positive eCO) or “non-smoker” (negative self-reported smoking verified by negative eCO, or self-reported status missing but negative eCO). Subjects without an eCO status were considered missing data.

Blood was collected at baseline for biomarker analysis and genotyping. A description of CC-16 measurement is given below. Genotyping in the ECLIPSE cohort was undertaken using the HumanHap 550 V3 (Illumina) and quality control was performed using BeadStudio, as previously described.[6, 7] The present analysis is based on the use ECLIPSE study data downloaded from the dbGaP web portal, under study accession #phs001252.v1.p1 (available: https://www.ncbi.nlm.nih.gov/projects/gap/cgi-bin/study.cgi?study_id=phs001252.v1.p1).

Human research ethics approval was granted by the institutional review boards at each of the participating centers.

**Serum CC-16 measurement in the studies**

Measurement of serum CC-16 levels in the LHS and ECLIPSE cohorts has been described previously.[8, 9] Briefly, blood samples were centrifuged and separated, and the serum frozen and stored until the time of analysis. CC-16 concentrations in thawed serum samples were measured using a commercially-available, sandwich enzyme linked immunosorbent assay (ELISA) (BioVendor, Heidelberg, Germany). Concentrations were determined by reference to a standard curve of known CC-16 concentrations.

In the LHS, subjects with serum CC-16 concentration below the lower limit of quantitation (LLQ) of 0.65 ng/mL were assigned a value equal to half the LLQ i.e 0.325 ng/mL.[9] For the present analysis, all subjects with CC-16 level equal to 0.325 ng/mL (n=240) were excluded due to their effect on the distribution of CC-16 concentrations. In ECLIPSE, subjects with serum CC-16 concentration below the LLQ were excluded; according to the study investigators, this equated to less than 1% of the cohort.[8] We further excluded outlier subjects from each cohort with serum CC-16 concentration >40 ng/mL (LHS, n=1, ECLIPSE, n=1). In order to better approximate a normal distribution, we transformed serum CC-16 concentrations by their natural logarithm prior their use in our analysis.

### 2. DATASET ANALYSED FOR ASSOCIATION WITH COPD: ICGC/UK BIOBANK META-ANALYSIS

A total of 35,735 COPD cases and 222,076 non-COPD controls within the International COPD Genetics Consortium (ICGC) and UK Biobank cohorts were included in a recent genome-wide association meta-analysis for the presence of COPD.[10]

#### Description of cohorts

The ICGC is an international collaboration of COPD cohort, case-control and general population studies with spirometry and genotype data available; the full description of the individual studies was previously described[11]. A case-control association analysis of the ICGC cohort was conducted based on prebronchodilator spirometry measurements, with COPD cases defined based on Global Initiative for Obstructive Lung Disease (GOLD) criteria.[12] Genotype imputation of the ICGC cohort was performed using the 1000 Genomes reference panel.[13] Human research approval for was obtained for each cohort in the ICGC, as previously described.[11]

The UK Biobank cohort is part of the UK Biobank project,[14] which involved deep phenotyping and genotyping of a total of 502,682 individuals. The majority of individuals within this cohort are white Europeans. COPD cases within the UK Biobank cohort were also defined according to GOLD criteria[12] based on prebronchodilator spirometry. Genotyping of the UK Biobank cohort was performed using the Affymetrix Axiom UK BiLEVE and UK Biobank array, and genotypes were imputed to the Haplotype Reference Consortium version 1.1 panel.[15] Human research ethics approval for the UK Biobank project was granted by the North West Multi-centre Research Ethics Committee (MREC). Oversight for the ethical conduct of the UK Biobank project is performed by the

UK Biobank Ethics Advisory Committee, for which the terms of reference can be found at https://www.ukbiobank.ac.uk/ethics/.

**Meta-analysis**

A logistic regression of COPD was conducted on the UK biobank data; this analysis was adjusted for sex, age, genotyping array, smoking exposure (pack-years), ever-smoking status, and genetic principal components.[10] The ICGC and UK biobank association results were combined by fixed-effect meta-analysis and statistically significant results were defined at the genome-wide significant threshold of $p<5x10^{-8}$. For our study, we used the resulting summary statistic of this meta-analysis.

### 3. GENOTYPE IMPUTATION AND QUALITY CONTROL

Genotypes in the LHS and ECLIPSE were imputed to the Haplotype Reference Consortium panel version 1.1 using the Michigan Imputation Server.[16] After imputation, we removed 218,380 SNPs with duplicated position information. For our GWASs, we only kept SNPs with minor allele frequency (MAF) > 0.01 and imputation quality ($r^2$) ≥ 0.7 (7,426,986 SNPs in LHS, and 7,471,599 SNPs in ECLIPSE). The GWASs assumed an additive genetic model adjusted for age, sex, BMI, smoking status (see section 1 – Description of study populations), and the first 5 genetic principal components. We then combined the results from the two cohorts using inverse variance weighted (IVW) fixed-effects meta-analysis. For the meta-analysis, we only included the 7,312,348 overlapping SNPs in both LHS and ECLIPSE.

In order to identify independently-associated SNPs, we performed conditional probability analysis[17] within each 2 Mb gene region using the Genome-wide Complex Trait Analysis (GCTA) platform version 1.92.0beta3[18] with the larger cohort (LHS) as the linkage disequilibrium (LD) reference. From this, we retained only those SNPs with independent, genome-wide significant ($p<5x10^{-8}$) association with CC-16 level.

### 4. MENDELIAN RANDOMISATION MODELS

Our Mendelian randomisation (MR) analysis relates the pQTL per-allele effects on serum CC-16 levels to their effects on COPD outcomes (COPD risk in the ICGC dataset, and change in FEV1 in the LHS/ECLIPSE). We used an inverse variance weighted MR model with the CC-16 pQTLs as instrumental variables (IVs), adjusting for LD between SNPs on the same chromosome calculated using PLINK v1.9[19, 20] on 503 European-descent samples from the 1000 Genomes Project Phase 3[13] and constraining the intercept to zero. MR analysis was performed using the MendelianRandomization v0.2.2 package in R.[21, 22]

MR analysis is only valid if the IVs meet a number of fundamental assumptions.[23] We employed several methods to test for violations of these assumptions. We used Cochran's Q test for heterogeneity, with significance set at a nominal p<0.05. The presence of significant heterogeneity suggests that the variability in the SNP associations is greater than would be expected by chance alone, and thus may indicate bias due to one or more invalid IVs.[24] Next, we performed MR-Egger analysis,[25] which is similar to the IVW model but with an unconstrained intercept term. This model therefore allows directional pleiotropy, the presence of which is suggested by a significant (i.e. non-zero) intercept term (nominal significance set at p<0.05). Finally, we performed the Mendelian Randomization Pleiotropy RESidual Sum and Outlier (MR-PRESSO) test in R.[26, 27] This test has three components: 1) a "global" test for directional pleiotropy, which compares the observed distance from the MR regression line (residual sum of squares) to the predicted distance from the regression line under the null-hypothesis of no directional pleiotropy; 2) the identification and removal of outliers that contribute to directional pleiotropy; and 3) a test for significant differences in the MR causal estimates before and after removal of outliers. Under the MR-PRESSO framework, components 2) and 3) only proceed if the global test for directional pleiotropy (1) is significant at a nominal p<0.05. A summary of the stregths and weaknesses of each of these methods for assessing IV validity is provided by Verbanck *et al*.[26]

### 5. DATASET ANALYSED FOR LUNG TISSUE GENE EXPRESSION: THE LUNG eQTL STUDY

**Description of cohorts**

The Lung eQTL Study[28] was a multi-national observational study examining non-tumor lung tissue samples collected from 1,111 people and 3 different sites (University of British Columbia (UBC), Laval University, and University of Groningen). The majority of samples were collected from study participants undergoing resection of presumed lung cancers. At the UBC site, n=39 samples were obtained at autopsy, n=22 from the diseased lungs of lung transplant recipients, and n=7 from donor lungs unsuitable for transplantation. For the majority of participants, lung function tests were performed immediately before surgery. 84% percent of the participants across all sites were current or ex-smokers.

Genotyping was performed on either blood (Laval site) or lung (UBC, Groningen sites) samples using the Illumina Human 1M-Duo BeadChip array, as previously described.[28]

Ethics approval for the Lung eQTL Study was granted by the Institut Universitaire de Cardiologie et de Pneumologie de Québec and the UBC-Providence Health Care Research Institute Ethics Boards. Lung tissue samples were collected according to the institutional review board guidelines for each the participating institutions. Written informed consent was obtained from all patients, or families of deceased patients where applicable.

### 6. ANALYSIS OF GENE EXPRESSION IN THE LUNG eQTL STUDY

Tissue sample processing in the Lung eQTL Study has been previously described.[28] Lung tissue gene expression (mRNA) was measured using the Affymetrix HI133 array consisting of 751 control probesets and 51,627 non-control probesets. Gene expression values were extracted using the Affymetric Power Tools software (Robust Multichip Average method).[29] After quality control filtering, normalized expression data were adjusted for age, sex and smoking status.

### 7. LUNG cis-eQTL DETERMINATION

We estimated the associations between SNP and mRNA expression by linear regression models assuming additive genotype effects in each site separately. We then performed a meta analysis to combine the results. We defined cis-eQTLs as within 1 Mb up- or downstream of the SNP.
