## Supplementary Results for "The protective effect of club cell secretory protein (CC-16) on COPD risk and progression: a Mendelian randomisation study"


**AUTHORS:**

Stephen Milne[1,2,3], Xuan Li[1], Ana I Hernandez Cordero[1], Chen Xi Yang[1], Michael Cho[4], Terri H Beaty[5], Ingo Ruczinski[6], Nadia N Hansel[7], Yohan Bossé[8], Corry-Anke Brandsma[9], Don D Sin[1,2], Ma'en Obeidat[1]

1. Centre for Heart Lung Innovation, St Paul's Hospital and University of British Columbia, Vancouver, BC, Canada
2. Division of Respiratory Medicine, Faculty of Medicine, University of British Columbia, Vancouver, BC, Canada
3. Faculty of Medicine and Health, University of Sydney, Sydney, New South Wales, Australia
4. Channing Division of Network Medicine and Division of Pulmonary and Critical Care Medicine, Brigham and Women's Hospital, Boston, MA, USA
5. Department of Epidemiology, Bloomberg School of Public Health, Johns Hopkins University, Baltimore, MD, USA
6. Department of Biostatistics, Bloomberg School of Public Health, Johns Hopkins University, Baltimore, MD, USA
7. Pulmonary and Critical Care Medicine, School of Medicine, Johns Hopkins University, Baltimore, MD, USA
8. Institut universitaire de cardiologie et de pneumologie de Québec, Department of Molecular Medicine, Laval University, Quebec City, Canada
9. University of Groningen Department of Pathology and Medical Biology, University Medical Centre Groningen, Groningen, The Netherlands

**CORRESPONDING AUTHOR:**

Dr Stephen Milne
UBC Centre for Heart Lung Innovation
Rm 166, St Paul's Hospital
1081 Burrard Street,
Vancouver, BC, V6Z 1Y6
CANADA

E:

T: +1 604 806 8346

1. **TABLE S1: Association between serum CC-16 level and change in $FEV_1$ in the biomarker cohorts**

| | n | Beta | SE | p |
|---|---|---|---|---|
| LHS | 3444 | 2.43 | 0.93 | 0.01* |
| ECLIPSE | 1821 | 6.25 | 3.94 | 0.11 |
| Meta-analysis | 5265 | 2.64 | 0.91 | 0.004** |

Multiple linear mixed effects model for change in $FEV_1$ with ln(CC-16), adjusted for age, sex, smoking status, baseline forced expiratory volume in 1 s (FEV1), body mass index, and their interaction with time. SE, standard error; LHS, Lung Health Study. *$p<0.05$ **$p<0.01$.

**2. TABLE S2: Associations between CC-16 protein quantitative trait loci (pQTLs) and COPD outcomes**

| | | | | COPD risk | | | COPD progression (Change in $FEV_1$ in mL/year) | | | | | |
|---|---|---|---|---|---|---|---|---|---|---|---|---|
| | | | | ICGC | | | LHS | | | ECLIPSE | | |
| SNP rsID | Chr | Position | Effect/Alt allele | Allele effect β | SE | p | Allele effect β | SE | p | Allele effect β | SE | p |
| rs4971100 | 1 | 155155731 | A/G | $-3.9x10^{-3}$ | 0.01 | 0.7 | -1.06 | 0.81 | 0.19 | 2.78 | 3.15 | 0.38 |
| rs1515498 | 3 | 189508302 | A/G | $-2.2x10^{-3}$ | 0.01 | 0.83 | 1.43 | 0.83 | 0.08 | 0.78 | 3.26 | 0.81 |
| rs37002 | 5 | 1356944 | C/T | -0.02 | 0.01 | 0.13 | -0.57 | 0.85 | 0.50 | -6.38 | 3.14 | 0.04* |
| rs11032840 | 11 | 34779464 | G/T | $3.8x10^{-3}$ | 0.01 | 0.71 | 0.10 | 0.79 | 0.90 | 1.49 | 3.08 | 0.63 |
| rs3741240 | 11 | 62186542 | G/A | -0.02 | 0.01 | 0.14 | 1.63 | 0.82 | 0.05* | 6.96 | 3.09 | 0.02* |
| rs11231085 | 11 | 62190448 | G/C | -0.02 | 0.01 | 0.11 | 2.89 | 0.83 | $5.4x10^{-4}$* | 4.79 | 3.28 | 0.14 |
| rs7962469 | 12 | 52348259 | A/G | -0.03 | 0.01 | $1.2x10^{-3}$* | -0.51 | 0.87 | 0.56 | -2.00 | 3.25 | 0.54 |

*p<0.05. ICGC, International COPD Genetics Consortium; COPD, chronic obstructive pulmonary disease; LHS, Lung Health Study; ECLIPSE, Evaluation of COPD Longitudinally to Identify Predictive Surrogate Endpoints; SNP, single nucleotide polymorphism; rsID, reference SNP cluster identifier; Chr, chromosome; Alt, alternate; SE, standard error.

**3. FIGURE S1: Manhattan plot for serum CC-16 genome-wide association study (GWAS) in Lung Health Study**

GWAS p values (–$\log_{10}$ scale) (Y axis) versus single nucleotide polymorphism positions across 22 chromosomes (X axis). Horizontal red line represents the genome-wide significance cut-off of $5x10^{-8}$. CC-16, club cell secretory protein-16.

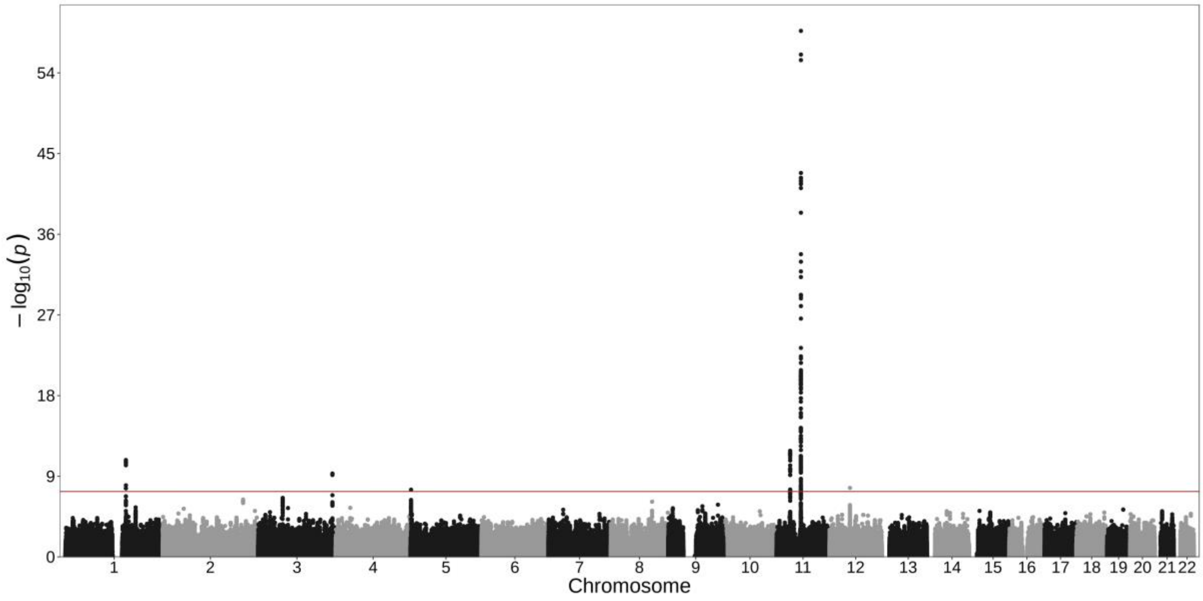

### 4. FIGURE S2: Manhattan plot for serum CC-16 genome-wide association study (GWAS) in ECLIPSE study

GWAS p values (–$\log_{10}$ scale) (Y axis) versus single nucleotide polymorphism positions across 22 chromosomes (X axis). Horizontal red line represents the genome-wide significance cut-off of $5x10^{-8}$. CC-16, club cell secretory protein-16.

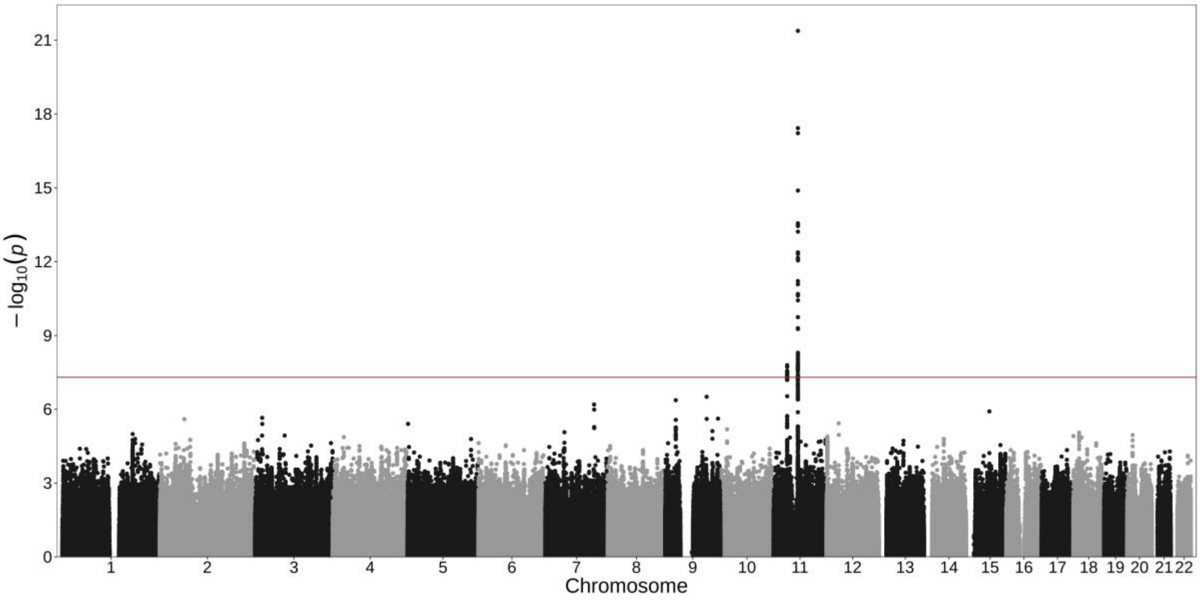

**5. FIGURE S3: Quantile-quantile plot for serum CC-16 genome-wide association study (GWAS)**

Meta-analysis of Lung Health Study and ECLIPSE study GWAS. CC-16, club cell secretory protein-16. Observed p-values (–$\log_{10}$ scale) (Y axis) versus expected p-values (–$\log_{10}$ scale) (X axis). Red line represents where observed p values are equal to the expected.

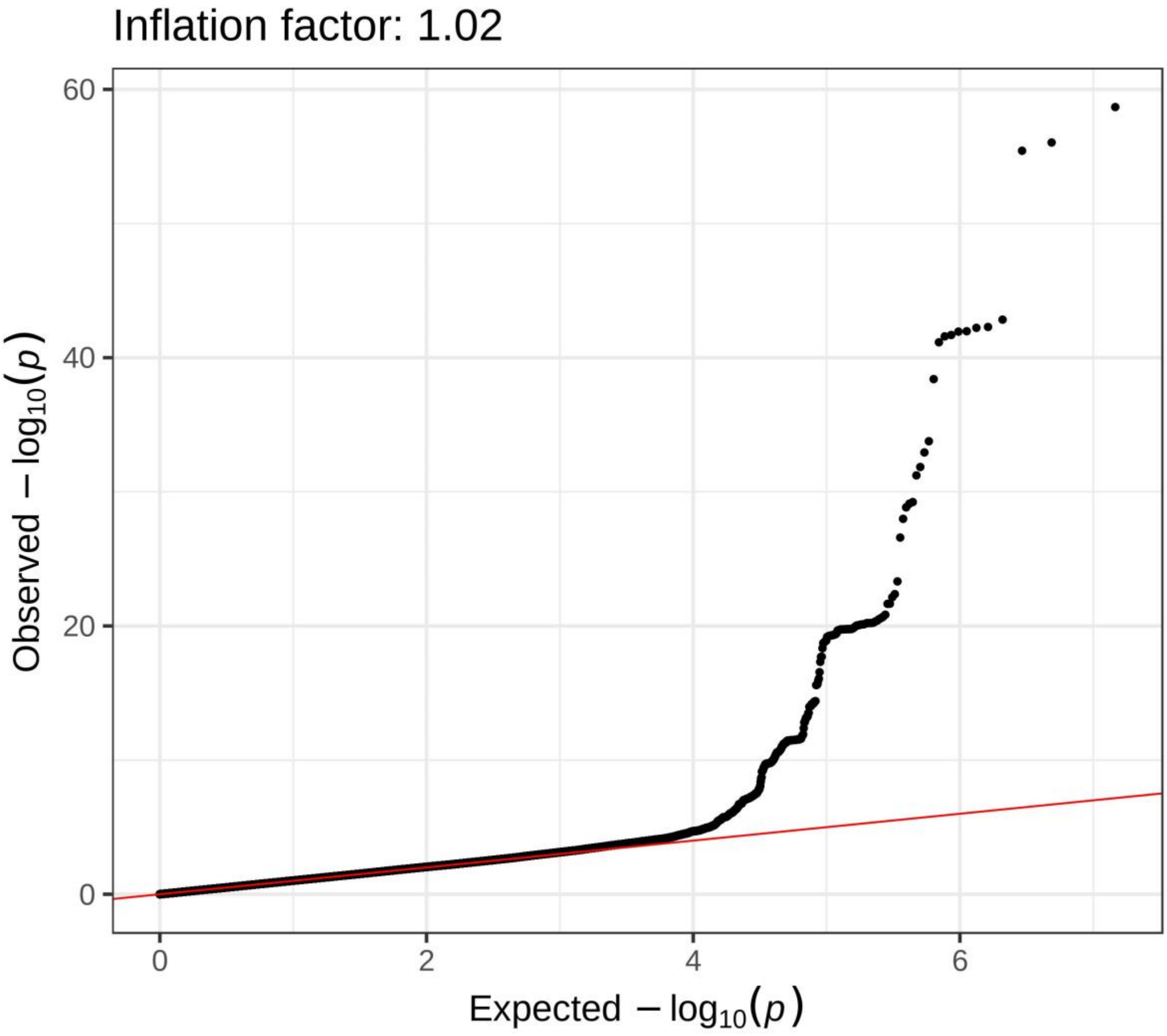

6. **FIGURE S4: Mendelian randomisation (MR) Egger analysis for COPD risk**

Inverse variance weighted regression model, adjusted for linkage disequilibrium between single-nucleotide polymorphisms (SNPs), with unconstrained intercept. The model relates the per-allele effects of the SNPs on serum CC-16 level to their per-allele effects on risk of chronic obstructive pulmonary disease (COPD) in the International COPD Genetics Consortium (ICGC) dataset. The red line represents the estimated effect. Error bars represent 95% confidence intervals. SNPs are annotated by their rs identifier. OR, odds ratio; CC-16, club cell secretory protein-16.

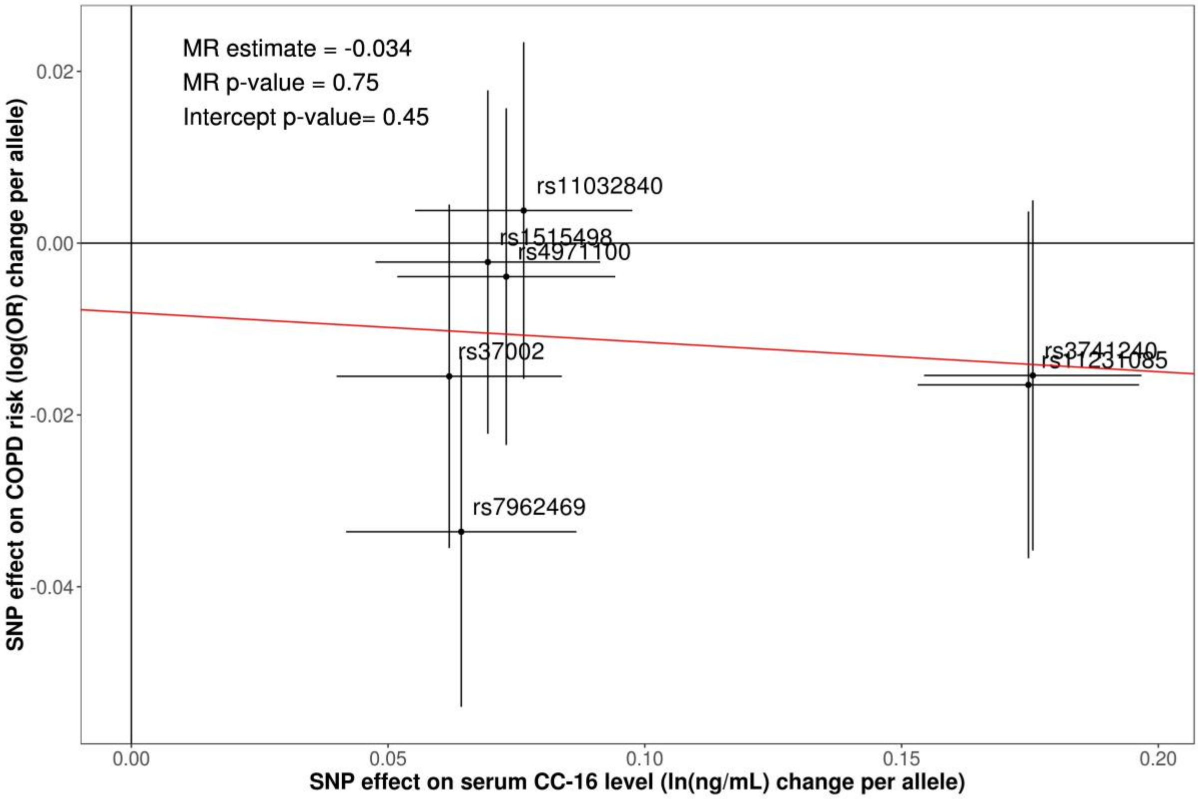

7. **FIGURE S5: Mendelian randomisation (MR) Egger analysis for COPD progression in the Lung Health Study**

Inverse variance weighted regression model, adjusted for linkage disequilibrium between single-nucleotide polymorphisms (SNPs), with unconstrained intercept. The model relates the per-allele effects of the SNPs on serum CC-16 level to their per-allele effects on lung function decline. The red line represents the estimated effect. Error bars represent 95% confidence intervals. SNPs are annotated by their rs identifier. CC-16, club cell secretory protein-16, $FEV_1$, forced expiratory volume in 1 second.

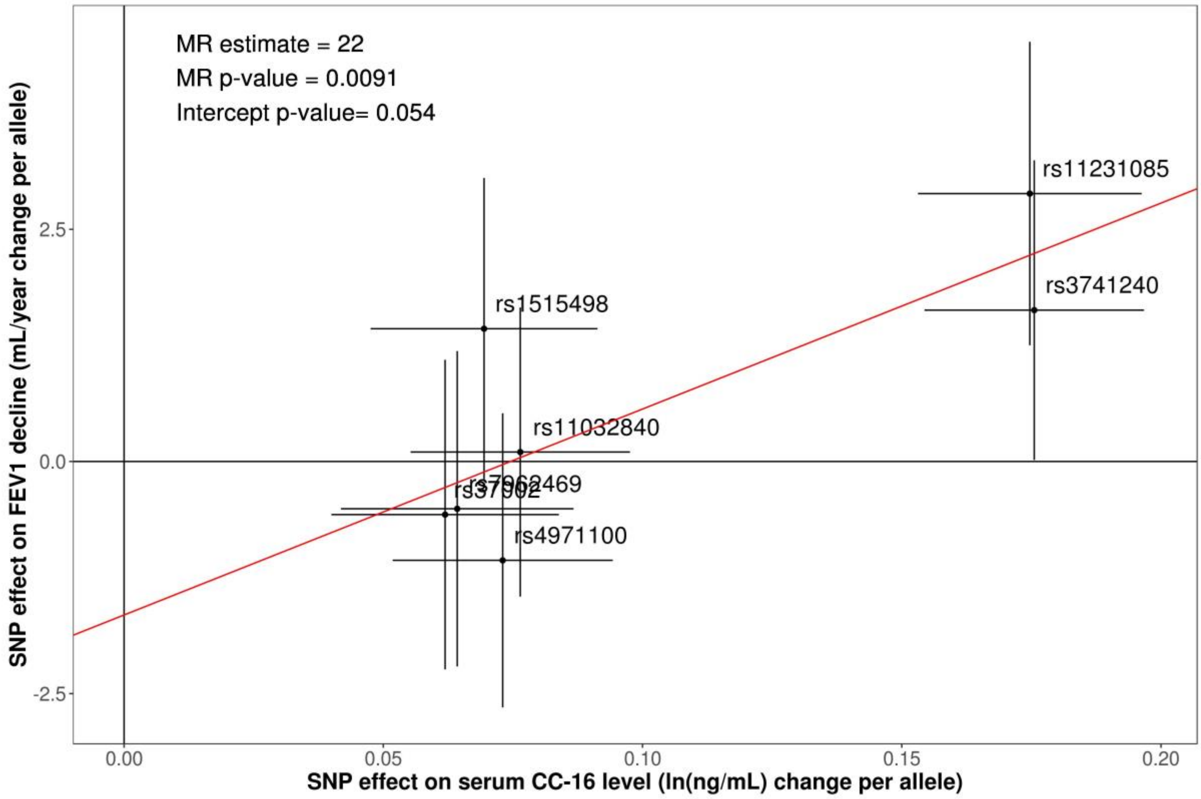

### 8. FIGURE S6: Mendelian randomization (MR) plot for COPD progression in ECLIPSE study

Inverse variance weighted regression model, adjusted for linkage disequilibrium between single-nucleotide polymorphisms (SNPs), intercept constrained to zero. The model relates the per-allele effects of the SNPs on serum CC-16 level to their per-allele effects on lung function decline. The red line represents the estimated effect. Error bars represent 95% confidence intervals. SNPs are annotated by their rs identifier. CC-16, club cell secretory protein-16; $FEV_1$, forced expiratory volume in 1 second.

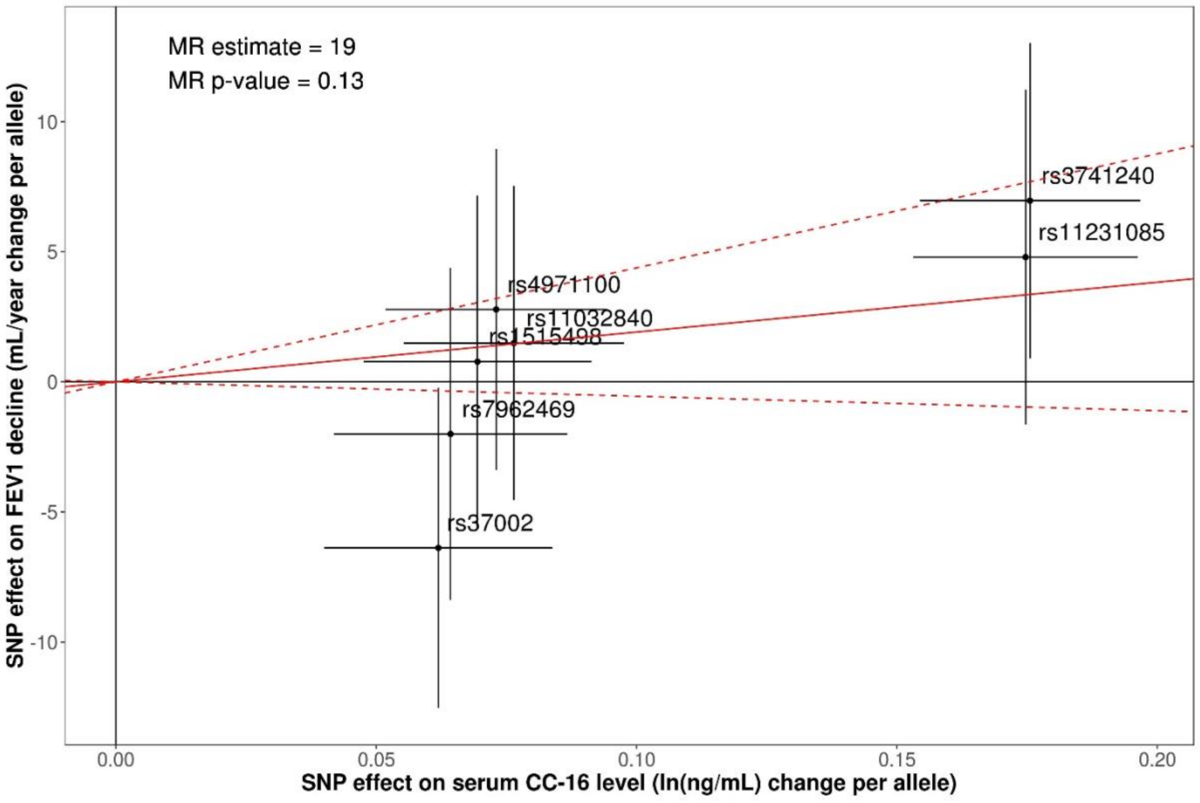
